# Morphology does not covary with predicted behavioural correlations of the domestication syndrome in dogs

**DOI:** 10.1101/660829

**Authors:** Christina Hansen Wheat, Wouter van der Bijl, Christopher West Wheat

## Abstract

Domesticated animals display suites of altered morphological, behavioural and physiological traits compared to their wild ancestors, a phenomenon known as the domestication syndrome (DS). Because these alterations are observed to co-occur across a wide range of present day domesticates, the traits within the DS are assumed to covary within species and a single developmental mechanism has been hypothesized to cause the observed co-occurrence. However, due to the lack of formal testing it is currently not well-resolved if the traits within DS actually covary. Here we test the hypothesis that the presence of the classic morphological domestication traits white pigmentation, floppy ears and curly tails predict the strength of behavioural correlations in support of the DS in 78 dog breeds. Contrary to the expectations of covariation among DS traits, we found that morphological traits did not covary among themselves, nor did they predict the strength of behavioural correlations among dog breeds. Further, the number of morphological traits in a breed did not predict the strength of behavioural correlations. Our results thus contrast with the hypothesis that the DS arises due to a shared underlying mechanism, but more importantly, questions if the morphological traits embedded in the DS are actual domestication traits or post-domestication improvement traits. For dogs, it seems highly likely that strong selection for breed specific morphological traits only happened recently in relation to breed formation. Present day dogs therefore have limited bearing of the initial selection pressures applied during domestication and we should reevaluate our expectations of the DS accordingly.

## Introduction

Domesticated animals display suites of altered morphological, behavioural and physiological traits compared to their wild ancestors, a phenomenon known as the domestication syndrome (DS). Key examples of components in the DS are increased tameness, reduced brain size, white pigmentation and decreased hypothalamic-pituitary-adrenal (HPA) axis reactivity (Kruska, 1996; Driscoll et al., 2009; Trut et al., 2009). Because these alterations are observed to co-occur across a wide range of present day domesticates, such as dogs (*Canis familiaris*), cats (*Felis catus*), rabbits (*Oryctolagus cuniculus*), horses (*Equus caballus)* and pigs (*Sus scrofa*) (Sánchez-Villagra et al., 2016), the traits within the DS are assumed to covary within species (Trut, 1998; Trut et al., 2009). Domestication experiments have demonstrated that selection for tame behaviour alone can produce the myriad changes seen in the DS (Belyaev et al., 1985; Trut et al., 2009). While the mechanistic origin of the DS in currently unresolved, these findings have nurtured the hypothesis that the convergent patterns seen across domesticated species arise via a singular developmental mechanism such as altered neuroendocrine control of ontogenesis (Belyaev, 1979), or neural crest deficit during embryogenesis (Wilkins et al., 2014). Both of these influential studies have led to the general assumption that morphological changes, such as white pigmentation, floppy ears and curly tails, have arisen as by-products of the physiological alterations caused by selection upon behaviour (Wilkins et al., 2014).

The hypotheses that the DS is founded in single developmental mechanism offer a coherent, logical and satisfying explanation for the observed covariation among DS traits. However, traits of the DS are not fully consistent with such hypotheses. First, DS traits are not evenly distributed among domesticated animals (Sánchez-Villagra et al., 2016). Second, even though rat (*Rattus norvegicus*) lines selected for tameness have an increased frequency of white spots (Trut et al., 2000), there is no genomic association between white coat colouration and tame behaviour (Albert et al., 2009). This is unexpected based on the hypothesis that white pigmentation should arise as a by-product of selection on tameness and further, because syndrome traits originating from a shared physiological origin should be difficult to decouple (*sensu* Sih et al., 2004). Finally, while recent genomic studies in horses (Librado et al., 2017), foxes (*Vulpes vulpes*, Wang et al., 2018), dogs (Pendleton et al., 2018) and cats (Montague et al., 2014) find signatures of domestication selection pressures in genes associated with neural crest development, and thus are argued to support the neural crest hypothesis, these genes are only a subset of many showing selective signatures during domestication. Thus, while it is generally assumed that DS traits covary, possibly due to a single developmental mechanism, further quantitative testing of this hypothesis is warranted.

Only recently has a formal test of covariance of DS traits been conducted. In their study of the behavioural component of the DS in more than 76,000 dogs, Hansen Wheat et al. (2019) demonstrated that while correlations between fear, aggression, sociability and playfulness were stronger in ancient breeds, these correlations were weaker or had been decoupled in modern breeds. However, this study focused only upon behaviour, which was likely the focal trait in dog domestication (*sensu* Belyaev et al., 1985; Trut et al., 2009). To date, no studies have investigated the covariation of morphological traits, either among themselves, or with the expected behavioural correlations of the DS. Such a formal investigation of the predicted expectations of how behavioural and morphological components of domestication arise is needed if we are to further our understanding of the DS.

Among domesticates, the dog has been argued to be the only species expressing the full DS (Sánchez-Villagra et al., 2016). Dogs have been bred for highly breed-specific morphological and behavioural traits (Svartberg, 2006; Mehrkam and Wynne, 2014), which are illustrated by the extreme phenotypic variation expressed among the more than 400 present day dog breeds (Lindblad-Toh et al., 2005; Parker et al., 2017). The result is dramatic phenotypic variation expressed across breeds. However, while key DS traits of behaviour and morphology do not qualitatively appear to occur simultaneously across breeds (Sánchez-Villagra et al., 2016), this has never been tested quantitatively. Furthermore, though dogs express a range of traits not present in wolves (Parker et al., 2009; Larson et al., 2014), it is currently not well resolved if dog traits are original domestication traits, i.e. traits evolved under direct selection in the initial stages of domestication, or so-called improvement traits that have been secondarily enhanced post-domestication during breed formation (Larson and Fuller, 2014; *sensu* Olsen and Wendel, 2013).

With modern breeds created from intense breeding efforts only within the last 150-200 years (Lindblad-Toh et al., 2005; vonholdt et al., 2010), it is possible that modern dogs provide a suboptimal basis for the expectations embedded in the DS. Indeed, as noted earlier, modern dogs lack the strong behavioural correlations expected of the DS (Hansen Wheat et al., 2019). Nonetheless, because the foundation for the DS hypothesis is based on extant domesticates, it remains unclear if we should expect the expression of the DS to vary across different stages of domestication. Archaeological findings of early dogs provide limited information on morphology (i.e. skeletal features), and none on behaviour, which impairs our ability to compare trait expression in dogs at different stages of domestication. Pre-breed formation domesticated dogs, i.e. village dogs, could be very informative, but unfortunately, the only non-admixed village dog populations identified to date are found in Borneo (Shannon et al., 2015) and have not been studied behaviourally. However, a small group of present day dogs can be categorized as ancient breeds due to their a) detectable admixture with wolf, which is not present in modern breeds, and b) an origin about 500 years ago (Lindblad-Toh et al., 2005; vonholdt et al., 2010). Certainly, ancient breeds are expected to have improvement traits, but importantly, these breeds have been shown to have stronger behavioural correlations expected of the DS compared to modern breeds (Hansen Wheat et al., 2019). While acknowledging they are an imperfect proxy, ancient breeds are arguably the only available representatives for earlier stages of dog domestication, and thus the division of ancient and modern breeds provides an opportunity for temporal comparisons among dogs on a domestication time scale.

Here we test the hypothesis that the presence of morphological traits predict the strength of behavioural correlations in support of the DS in dogs. For the morphological component of our study, we focused upon variation in the traits white pigmentation, floppy ears and curly tails, which have been referred to as morphological markers of domestication (Trut et al., 2009). For the behavioural component, we used estimates of effect sizes for behavioural correlations associated with the DS, derived from data extracted from the Swedish Kennel Club’s database on 76,158 dogs completing a highly standardized behavioural test battery (Hansen Wheat et al., 2019). We then matched these effect sizes of behavioural correlations with our estimates of morphological traits from the 78 breeds. We further added a temporal component by assessing 7 ancient and 71 modern breeds separately. As predicted by the DS, we expected that the presence of white pigmentation, floppy ears and curly tails would co-vary among breeds. Additionally, we expected that the presence or absence of these morphological traits would predict the strength of behavioural correlations of the DS. That is, we expected stronger behavioural correlations of the DS when morphological traits of the DS are present, as well as the converse, weaker behavioural correlations when morphological traits of the DS are absent. We further predicted that behavioural correlations would be stronger with the number of morphological traits present.

## Methods

### Morphological assessment

We based our study on the 78 dog breeds used in a recent study to test behavioural correlations within the domestication syndrome (Hansen Wheat et al., 2019). Of the 78 breeds, seven were ancient breeds and 71 modern breeds. This difference in sample sizes between breed groups does not reflect a lack in sampling effort, but the natural limitation of only few breeds being categorized as ancient. We carefully inspected the breed standards for those 78 breeds by consulting the Fédèration Cynologique Internationale, the world’s largest federation of kennel clubs, to assess the presence or absence of our three chosen morphological traits; white pigmentation, floppy ears and curly tails. We used both relaxed and conservative assessments of the three morphological traits (Figure 1, Figure 2, Figure S1). We defined white pigmentation as any form of white pigmentation in the breed, regardless of its placement or shape. We also classified dogs with a white base colour, such as Dalmatians and Samoyeds, to express white pigmentation. Breeds where “white” was not mentioned in the coat colour description, such as Dobermann and Rottweiler, were assessed as not having white pigmentation. For our conservative assessment of white pigmentation, only breeds specifically described to have a white base colour or characteristic white coloration, or breeds where some versions have white pigmentation (such as Schnauzers) were included. Breeds were a small white spot or a few white hairs are “tolerated” or “undesirable” were not included as having white pigmentation in our conservative assessment. For the relaxed assessment, we included breeds were small white spots or a few white hairs, for instance on the chest, are “tolerated” or “undesirable” (Figure 1A-D). Floppy ears were assessed based on whether a breed has ears that are either erect or to some degree floppy (i.e. from just the tip to hanging straight down, Figure 1E-H). Thereby the presence or absence of floppy ears was assessed as a completely binary trait, and did not differ between the relaxed and conservative assessments. For our conservative assessment of curly tails, only breeds described to specifically have their tail in a permanent curl, i.e. with no option to let down the tail, as seen in Pugs, were included. For the relaxed assessment breeds that are described to carry their tail in a “curl”, “hook”, “sabre”, “sickle” or “J”, and even breeds carrying their tails in the slightest “curve”, but can let their tails straight down were assessed as having curly tails (Figure 1I-L). Breeds where the words “curl”, “hook”, “sabre”, “sickle”, “J” or “curve” were not included in the description of the tail were assessed as not having a curly tail in either assessment.

**Figure 1.**
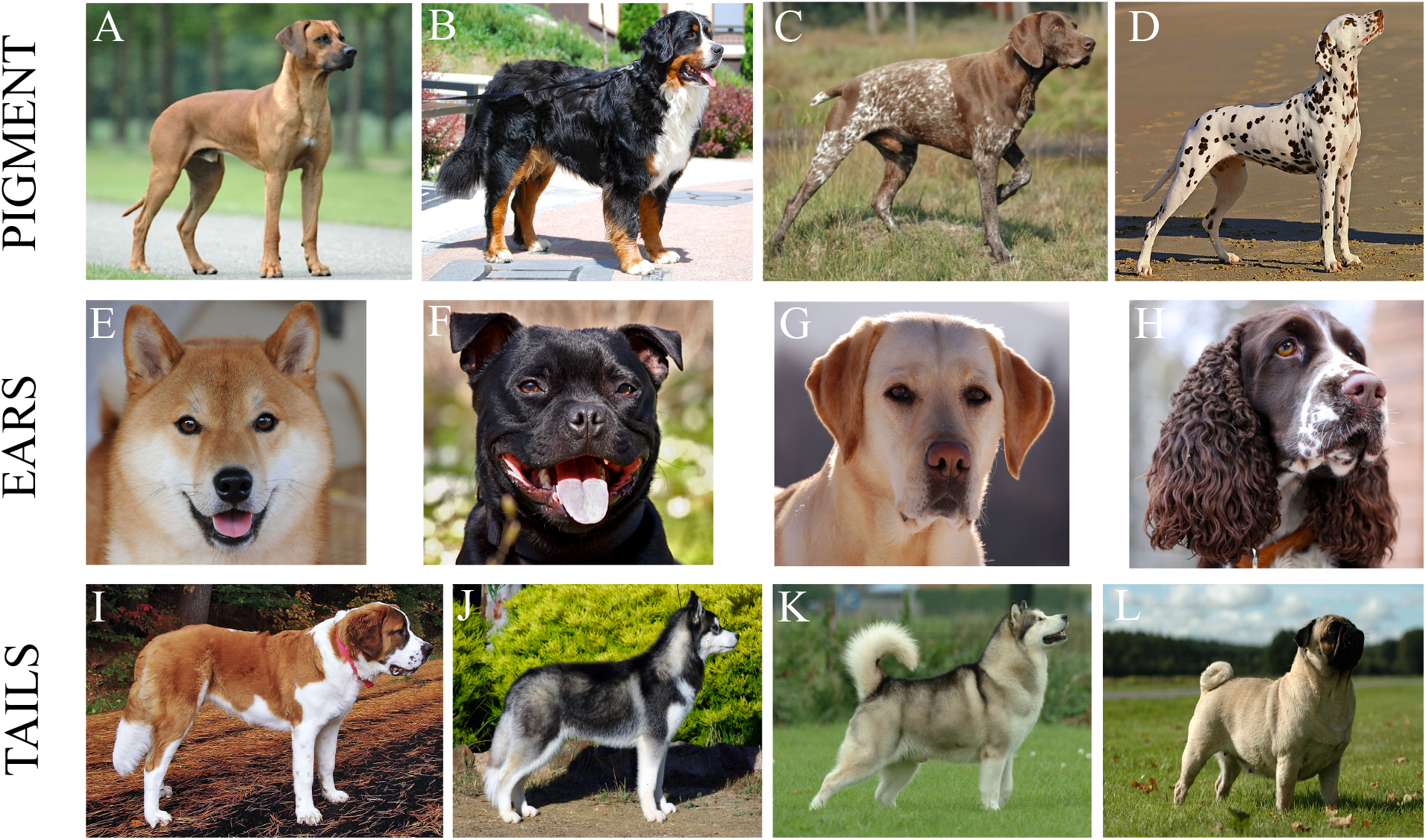
Morphological assessments. Examples of morphological variation across dog breeds and how this was taken into account when assessing the presence and absence of morphological traits in the DS. ***White pigmentation (pigment, A-D):*** Breeds where a small white spot or a few white hairs on the chest is tolerated or undesirable, here illustrated in a Rhodesian Ridgeback (A), were categorized as not having white pigmentation in the conservative assessment, but as having white pigmentation in the relaxed assessment. The presence of white pigmentation varies across breeds in size, shape and placement as illustrated in Bernese Mountain Dog (B), German Short-haired Pointer (C) and Dalmatian (D). ***Floppy ears (ears, E-H):*** Floppiness of ears is binary and erect ears, as illustrated in the Shiba (E), can never be floppy. Other examples of breeds with erect ears are Siberian Husky (J) and Alaskan Malamute (K). The floppiness of ears can be graduated as illustrated by the Staffordshire Bull Terrier (F), Labrador Retriever (G) and English Springer Spaniel (H). Any degree of floppiness of the ears was assessed as presence of floppy ears. ***Curly tail (tails, I-L):*** Breeds, such as the St. Bernard (I), with tails hanging straight down and never carry their tail in a curl, curve, hook, sickle, sabre of J shaped express the absence of a curly tail (both assessments). Many breeds carry their tail in a curl, curve, hook, sickle, sabre of J shaped fashion, but can also let their tail straight down, here illustrated by Siberian Husky with a let down tail (J) and an Alaskan Malamute with a tail carried in a curl (K). For the conservative assessment such breeds were categorized as not having curly tails, while they were categorized as having curly tails in the relaxed assessment. Other examples of breeds categorized like this are Rhodesian Ridgeback (A) and Dalmatian (D). A few breeds, like Pugs (L), express the presence of a permanent curly tail (both assessments). All photos are from wikicommons, please see references for specific credits.

### Behavioural assessment

For the behavioural component of our study, we used the dataset presented in Hansen Wheat et al. 2019, in which the strength and direction of behavioural correlations between aggression, fearfulness, sociability and playfulness across the 78 dog breeds were investigated. Behavioural data were provided by the Swedish Kennel Club for dog completing the Dog Mentality Assessment, a highly standardized behavioural test for dogs in Sweden. We refer to this paper for a full description of the methods used to estimate the effect sizes for these behavioural correlations.

### Statistical analyses

To evaluate the relationship between breed morphology and agreement with the domestication syndrome hypothesis, we assessed the correlation between our morphology scores, treated as dichotomous variables. First, we estimated the phi coefficient (*φ)* for presence/absence of each of each trait in pairwise combinations with significance determined using Fisher’s Exact Test, as implemented in the xtab_statistics function of the sjstats package v. 0.17.5 (Lüdeke, 2019). Second, we repeated this analysis using a Pearson’s product-moment correlation with similar results. Third, we assess whether the presence/absence of traits were correlated while taking into account phylogenetic correction, using a pairwise bionomial phylogenetic glm.

To evaluate the relationship between breed morphology and agreement with the domestication syndrome hypothesis, as quantified by the strength and direction of behavioural correlations, we used a meta-analytic model. It is a multi-level model that uses the 1326 observed correlation coefficients (Hansen Wheat et al., 2019), and their associated uncertainty, as the dependent variable. These correlations test multiple behavioural predictions by the DS, such as a positive association between sociability and playfulness, or a negative association between sociability and aggression (Trut et al., 2009; Himmler et al., 2013). The correlations test six such DS predictions. For some predictions multiple correlations per breed were measured, since the Dog Mental Assessment test provided multiple measurements for aggression and fearfulness. 17 correlations were obtained per breed. Therefore, we treat the DS as a nested compound hypothesis, with six predicted associations and 17 correlations. We aligned the sign of the correlations with the predicted directions, i.e. we flipped the sign of correlations expected to be negative, so that positive effect sizes represent support in favour of the DS.

To account for this nested structure, we included group level effects that allow the support for the DS to vary between the different predicted associations and the measured correlations. We additionally included group level effects of morphology for the associations and correlations, so that the moderating effect of morphological traits could be stronger or weaker depending on what behavioural correlations were measured. Since each breed was represented by multiple correlations, we included a group level intercept for breed. And because breeds are nonindependent due to shared ancestry (Felsenstein, 1985), an additional group level effect was added with the expected covariance matrix of the phylogeny. Morphology was modelled as three additive binary effects, one each for the presence or absence of white pigmentation, floppy ears and curly tails. We implemented the models in the probabilistic programming language Stan (Carpenter et al., 2017), using the interfacing R (R Core Team, 2019) package brms (Bürkner 2017, 2018). In brms syntax, the models were of the form: *Zr* | *se*(*vi, sigma = TRUE*) ~ *breed_category + pigmentation + ears + tails* + (1 + *breed_category* + *pigmentation* + *ears* + *tails* || *prediction/correlation*) + (1 | *breed*) + (1 *phylogeny*), where Zr is the z-transformed correlations coefficients, vi is the measurement error and sigma = TRUE allows for the estimation of the residual standard deviation.

Inference about the effects of morphology was based on two approaches. We used the posterior distributions for the parameters directly to evaluate the role of the three morphological traits separately. Secondly, we assessed the role of the number of morphological traits (regardless of which trait) by calculating the estimated mean response for each trait combination, and then calculating the marginal mean for a breed having 0, 1, 2 or 3 traits present.

Posterior distributions for the parameters were obtained through MCMC sampling, using 16 chains of 2000 iterations each, of which 1000 were warmup. We adjusted the adapt delta to 0.995 and the maximum tree depth to 20 to eliminate any divergent transitions. For population level effects, we used the default weak student-t prior with a mean of 0, scale parameter of 10 and 3 degrees of freedom. The same prior was used for standard deviations of group-level effects and the residual standard deviation, but there it was restricted to be non-negative. Trace plots indicated that the chains were well mixed, and we obtained an effective sample size of more than 2500 for all parameters. The largest *Ȓ* was 1.01, indicating convergence.

All analyses were done for both relaxed and conservative assessments of morphological traits. Results for the two different assessments were qualitatively similar, and below we present the results for the relaxed assessment (see Supplemental Files for results for the conservative assessment)

## Results

We placed the morphological traits and average effect sizes for behavioural correlations onto the latest dog phylogeny (Parker et al., 2017), revealing large variation among breeds in both our morphological and behavioural traits (Figure 2, for conservative assessments see Supplemental Files and Figure S1).

**Figure 2.**
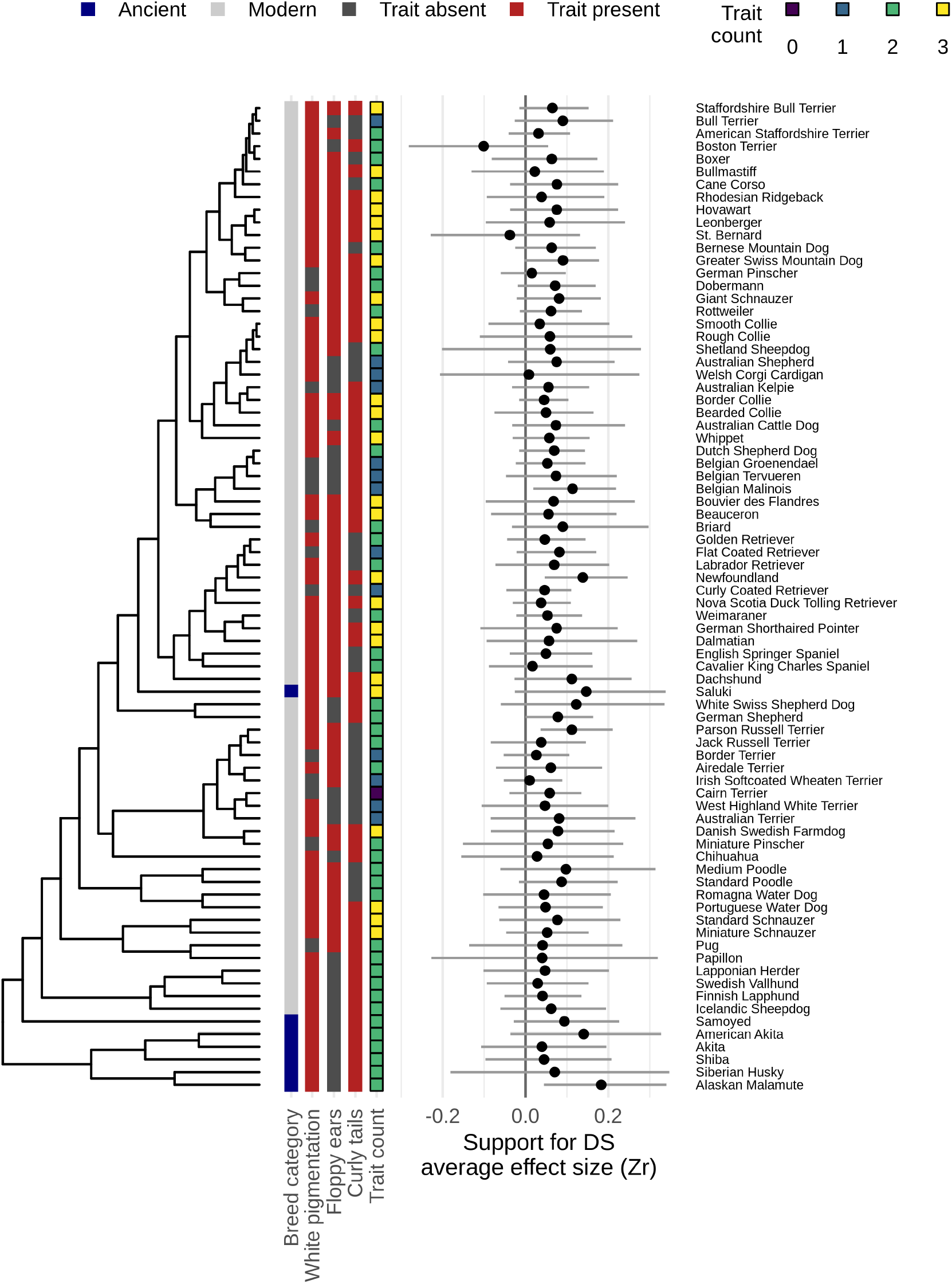
Morphological scores placed onto the latest dog phylogeny. Morphological scores based on the presence or absence of curly tail, floppy ears and white pigmentation (relaxed assessment), and average effect sizes for behavioural correlations in ancient and modern dog breeds placed onto the latest dog phylogeny (Parker et al 2017). Average effect sizes were calculated by separate meta-analytic models per breed (not used for inference), and posterior means + −95% credible intervals are depicted.

First, we used three different methods to test whether the presence of morphological DS traits covary amongst themselves. Neither phi coefficients (φ), Pearson’s product-moment correlation (t) nor phylogenetically corrected correlations (z) for the three morphological traits produced significant results: white pigmentation vs. floppy ears (φ = 0.172, p_φ_ = 1; t = − 0.115, p_t_ = 0.909; z = −0.080, p_z_ = 0.937), white pigmentation vs. curly tail (φ = 0.013, p_φ_ = 1; t = −5.7071^−20^, p_t_ = 1; z = 0.653, p_z_ = 0.514), floppy ears vs. curly tail (φ = 0, p_φ_ = 0.2063; t = −1.5176, p_t_ = 0.1333; z = −0.49807, p_z_ = 0.618).

Second, to test whether the presence of white pigmentation, floppy ears and curly tails predicts the strength of any of the behavioural correlations, we evaluated these traits as binary predictors of DS support. We found that there was no difference in the behavioural correlations when any of the three morphological traits were present or absent (Table 1, Figure 3A and B, Figure S2 and Supplemental Files). We emphasize that there is no support for even a very small difference in effect size (most extreme effect within CI: 0.04, Table 1). We did not confirm an effect of breed age, as the difference between ancient and modern breeds could not be clearly distinguished from 0, although considerable uncertainty in this estimate remains and most of the posterior favours stronger behavioural correlations in ancient breeds (Figure 3A and B, Table 1, for non conservative measurements see Supplemental Files).

**Figure 3.**
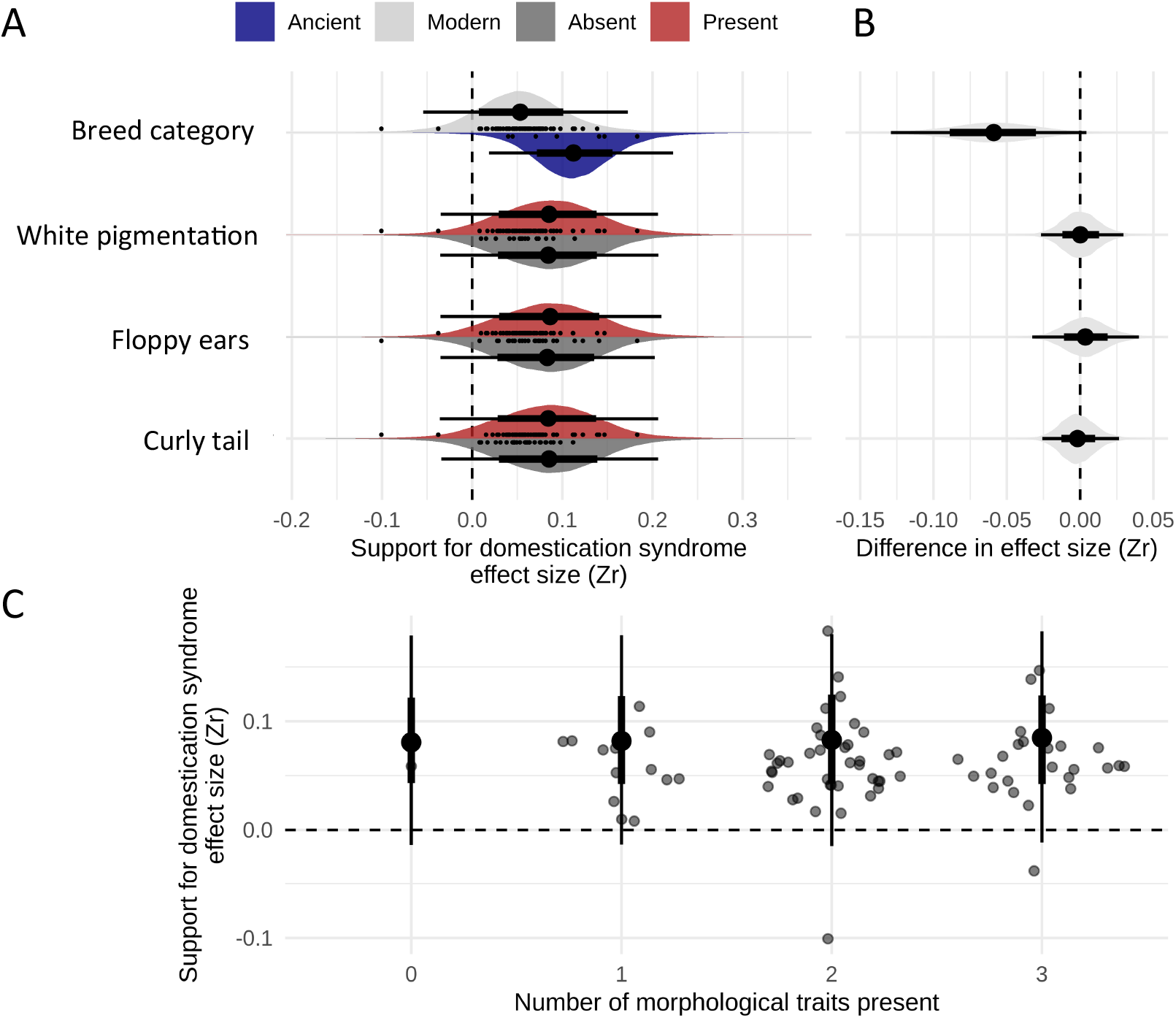
Morphological traits and the strength of behavioural correlations. A) Estimated support for the DS, quantified as the strength of behavioural correlations (*Z_r_*) depending on the presence or absence of morphological traits (relaxed assessment) and trait category. B) Regression coefficients indicating the difference between binary categories, as in A). C) The number a morphological traits present (relaxed assessment), i.e. morphological score, related to the estimated strength of behavioural correlations within the DS. In all panels, density distributions depict the full posterior distributions, with the thick lines covering the 66% credible interval, thin lines the 95% credible interval and point estimate the posterior median. Scattered points in A) and C) are the estimated average effect size per breed (as in Figure 2).

**Table 1.** Predictive value of morphological traits. Predictive value of the presence or absence of morphological traits on the strength of behavioural correlations in the DS. Posterior mean, posterior standard deviation (sd) and 95% Credible Interval (CI) given for breed category (ancient and modern) and the three morphological traits white pigmentation, floppy ears and curly tail.

| Term | Posterior mean | Posterior sd | 95CI<sub>lower</sub> | 95CI<sub>upper</sub> |
| --- | --- | --- | --- | --- |
| **Intercept** | 0.113 | 0.053 | 0.02 | 0.22 |
| **Breed category** | -0.06 | 0.034 | -0.129 | 0.004 |
| **White pigmentation** | 0.001 | 0.014 | -0.027 | 0.03 |
| **Floppy ears** | 0.004 | 0.019 | -0.033 | 0.04 |
| **Curly tail** | -0.001 | 0.014 | -0.026 | 0.026 |

Lastly, we evaluated support for the DS based on the “morphology score” of each breed, which ranged from 0 − 3 depending on how many, if any, of the three morphological traits is present in a breed (Figure 3C, Supplemental Files). We found that the number of morphological traits present in a breed did not predict the strength of behavioural correlations (0 traits: posterior mean_slope_ = 0.080, 95CI[-0.006 − 0.191]; 1 trait: posterior mean_slope_ = 0.080, 95CI[-0.008 − 0.193]; 2 traits: posterior mean_slope_ = 0.082, 95CI[-0.006 − 0.192]; 3 traits: posterior mean_slope_ = 0.083, 95CI[-0.004 − 0.190]). Given the small number of ancient breeds, we were not able to include breed age in this morphology score analysis.

## Discussion

Here we tested the whether the presence of three traits referred to as the morphological markers of domestication (white pigmentation, floppy ears and curly tails) predicted the strength of behavioural correlations within the DS. Contrary to the expectations of covariation among DS traits, we found that these morphological traits did not covary among themselves, nor did they predict the strength of behavioural correlations among dog breeds. Further, the number of morphological traits in a breed did not predict the strength of behavioural correlations. Additionally, we found no effect of breed age, i.e. ancient and modern breeds, in the predictive value of morphological traits on behavioural correlations. A high covariance among DS traits suggests a strong, central role for their shared origin in a single developmental source (e.g. white pigmentation arising as a by-product of increased tameness, Wilkins et al., 2014), while a lack of covariance suggests a more complex genotype to phenotype relationship. Thus, the lack of covariation among morphological and behavioural traits in our study is not consistent with the hypothesis that trait alterations in the DS are founded in a singular developmental source (Belyaev, 1979; Wilkins et al., 2014).

The DS in animals is primarily based on observations in present day domesticates. However, the ability of phenotypes in extant domesticates to provide insights about selection during initial domestication is complicated by post-domestication selection events, i.e. improvement traits (Olsen and Wendel, 2013; Larson and Fuller, 2014). Initial domestication efforts likely targeted existing variation at multiple loci across the genome (Larson and Fuller, 2014), but the breed-specific morphology and behaviour expressed in present day dog breeds was likely selected for post-domestication during breed formation. Many of the morphological traits seen across modern dog breeds are therefore not likely to be by-products of initial selection for domestication traits rather they are most likely improvement traits. Thus, while studies refer to the phenotypes of modern dog breeds as evidence for the DS (Wilkins et al., 2014; Sánchez-Villagra et al., 2016), whether these traits are relevant to domestication itself is questionable. Thus, our findings of a lack of covariation among morphological and behavioural traits, rather than providing insights into the DS, could be due to these traits being improvement traits, for which no covariance is expected. Regardless, the phenotypes of modern dog breeds should be interpreted with caution when trying to understand the domestication process.

One way to gain more insight into selection pressures during earlier stages of dog domestication, rather than those of breed improvement, is to include a temporal comparison by separating out ancient breeds and modern breeds. Here, we investigated whether the presence of morphological traits predict the strength of behavioural correlations in each breed group, but found no such effect. This finding contrasts with a recent study in which behavioural correlations of the DS were demonstrated to be stronger in ancient breeds compared to modern breeds (Hansen Wheat et al., 2019). Given that selection on tameness alone can generate the DS in foxes (Trut et al., 2009), and that aggression shows selective signatures directly associated with initial domestication efforts in these selection lines of foxes (Kukekova et al., 2018), it is likely that initial selection during dog domestication acted upon behaviour, not morphology (*sensu* Belyaev et al., 1985; Trut et al., 2009). Thus, with behaviours in the DS likely representing domestication traits, behavioural domestication phenotypes might to a larger extent be maintained in ancient compared to modern breeds. Morphology in dog breeds on the other hand, is arguably linked to breed improvement (Larson and Fuller, 2014), as reflected in the large variability in morphological trait combinations across dog breeds as quantified here.

In sum, whether the lack of covariance between morphology and behaviour in dogs is due to decoupling of independent domestication alleles (possibly caused by altered selection regimes during breed formation), these traits never having covaried because of a singular developmental mechanism or whether it is because we are applying a domestication hypothesis on traits that are not actual domestication traits, but rather improvement traits, remains an open question. If the latter is true, which seems likely for dogs, we must reevaluate our expectations of the DS and thereby also our assessment of DS traits in present day domesticates, as they have limited bearing of the initial selection pressures applied during domestication.

## Author contributions

CHW and CWW conceived the study. CHW prepared the data and all authors discussed how to analyse it. WvdB and CWW analysed the data. CHW prepared the manuscript draft and WvdB and CWW provided comments to produce the final version.

## Data availability statement

All data used for this study is available through Figure 2, the Supplemental Materials and Hansen Wheat et al. 2019.

