## Supplemental Figures for "Morphology does not covary with predicted behavioural correlations of the domestication syndrome in dogs"

**Figure S1.** Morphological scores on dog phylogeny (conservative) ……………………. 2

**Figure S2.** Predictive value of morphological traits (relaxed) ………………………… 3

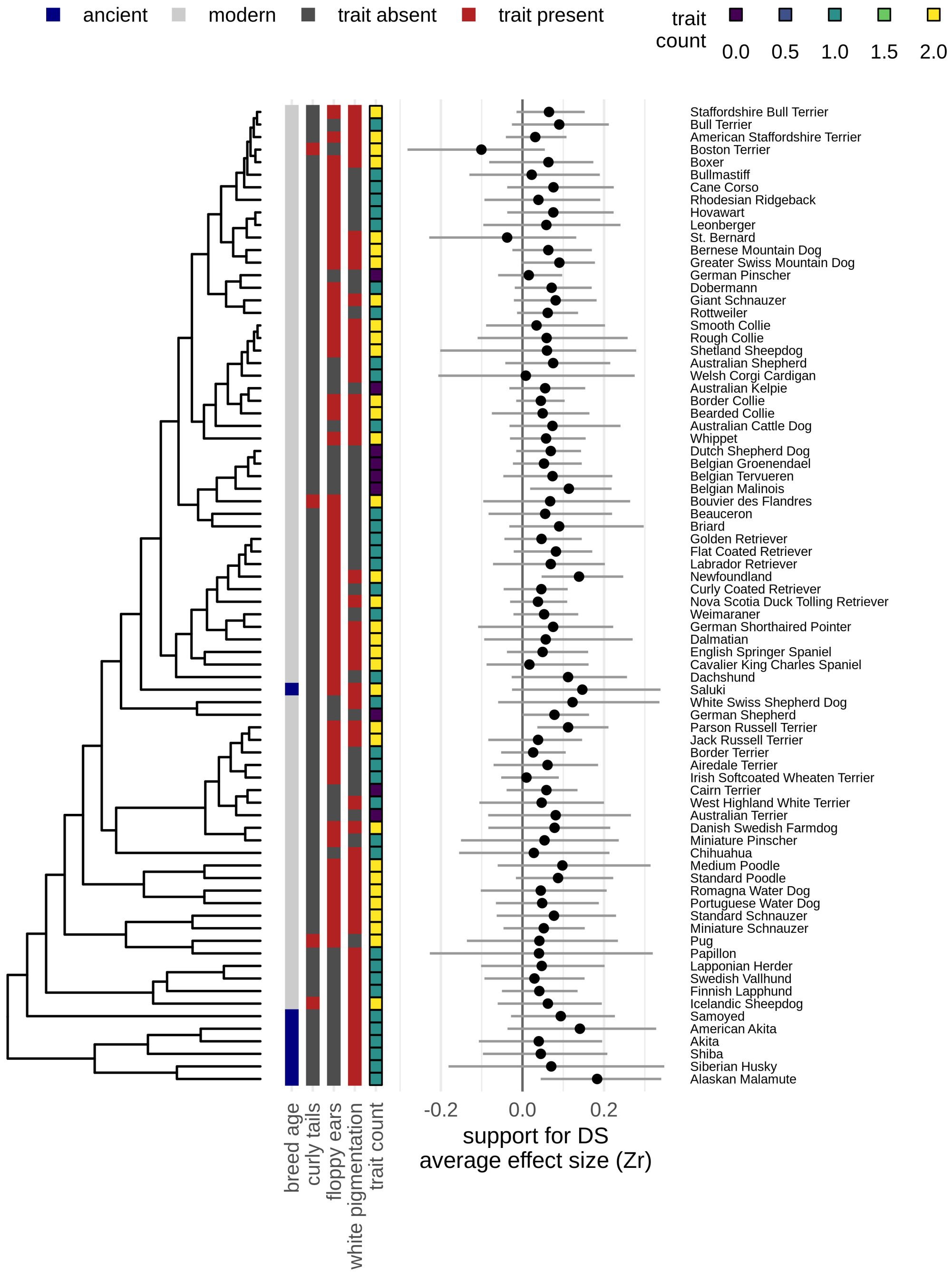


**Figure S1. Morphological scores on dog phylogeny (conservative).** Morphological scores based on the presence or absence of curly tail, floppy ears and white pigmentation (relaxed assessment), and average effect sizes for behavioural correlations in ancient and modern dog breeds placed onto the latest dog phylogeny (Parker et al 2017). Average effect sizes were calculated by separate meta-analytic models per breed (not used for inference), and posterior means +- 95% credible intervals are depicted.

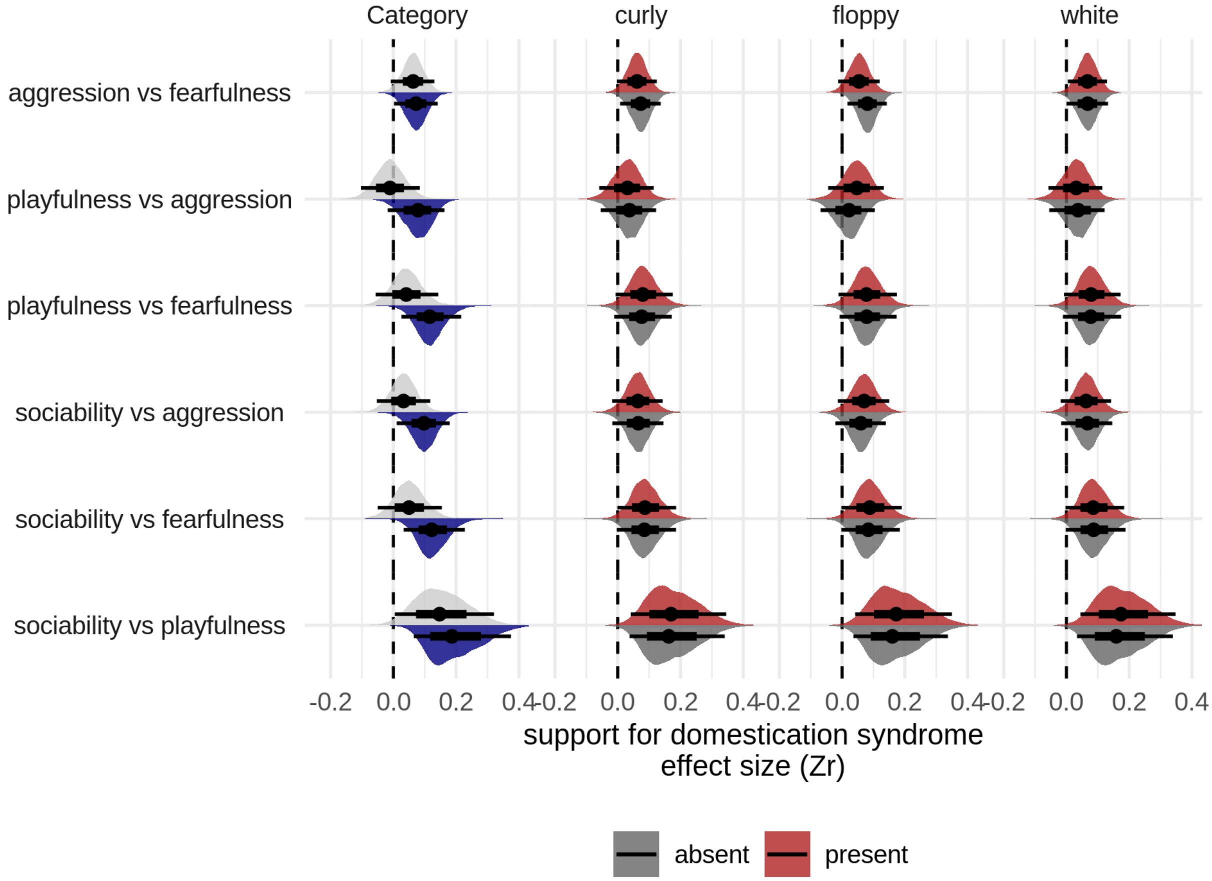


**Figure S2. Predictive value of morphological traits (relaxed).** The predictive value of the presence or absence of white pigmentation, floppy ears and curly tail on the support for the DS, quantified as the posterior distribution of the strength of the six individual behavioural correlations ($Z_r$) aggression vs fearfulness, playfulness vs aggression, playfulness vs fearfulness, sociability vs aggression, sociability vs fearfulness and sociability vs playfulness. None of the distributions are significant.
